# Frontal-Auditory Cortical Interactions and Sensory Prediction During Vocal Production in Marmoset Monkeys

**DOI:** 10.1101/2024.01.28.577656

**Authors:** Joji Tsunada, Steven J. Eliades

## Abstract

The control of speech and vocal production involves the calculation of error between the intended vocal output and the resulting auditory feedback. Consistent with this model, recent evidence has demonstrated that the auditory cortex is suppressed immediately before and during vocal production, yet is still sensitive to differences between vocal output and altered auditory feedback. This suppression has been suggested to be the result of top-down signals containing information about the intended vocal output, potentially originating from motor or other frontal cortical areas. However, whether such frontal areas are the source of suppressive and predictive signaling to the auditory cortex during vocalization is unknown. Here, we simultaneously recorded neural activity from both the auditory and frontal cortices of marmoset monkeys while they produced self-initiated vocalizations. We found increases in neural activity in both brain areas preceding the onset of vocal production, notably changes in both multi-unit activity and local field potential theta-band power. Connectivity analysis using Granger causality demonstrated that frontal cortex sends directed signaling to the auditory cortex during this pre-vocal period. Importantly, this pre-vocal activity predicted both vocalization-induced suppression of the auditory cortex as well as the acoustics of subsequent vocalizations. These results suggest that frontal cortical areas communicate with the auditory cortex preceding vocal production, with frontal-auditory signals that may reflect the transmission of sensory prediction information. This interaction between frontal and auditory cortices may contribute to mechanisms that calculate errors between intended and actual vocal outputs during vocal communication.

## Introduction

Our ability to control vocal production is a result of self-monitoring mechanisms that calculate and correct errors between intended and actual vocal outputs.^1–4^ This vocal control is important for efficient communications with other individuals, ensuring accurate vocal production, an ability seen not only for humans but also for many animal species. For example, both humans and non-human primates exhibit control of their vocal amplitude that affected by the presence of environmental noise, known as the Lombard effect.^5–9^ Similar control is seen for other aspects of vocal acoustics, including experimental manipulations using frequency-shifted auditory feedback that leads to behavioral compensation in humans and monkeys.^1,10,11^

Recent evidence has demonstrated potential neural correlates for such vocal error calculation in the auditory cortex. During vocal production, there is a well-described suppression of the auditory cortex that begins prior to the onset of vocalization.^12–20^ However, despite this suppression, these same auditory cortical neurons exhibit sensitivity to differences between actual vocal output and experimentally-altered auditory feedback^11,21–23^, suggesting that the suppression of the auditory cortex may reflect an error calculation. Importantly, there is also a correlation between feedback-sensitive activity in the auditory cortex and behavioral compensation to altered vocal feedback that implies a role in feedback-dependent vocal control.^11,24^ Because error calculation requires information not only about vocal sensory feedback but also about intended vocal outputs^3,4^, the auditory cortex has been hypothesized to receive top-down sensory predictions from brain structures that initiate and control vocal production. These predictions, termed efference copies or corollary discharges, are common to many sensory-motor systems.^25–30^ However, the underlying mechanisms and neural circuits remain uncertain.

One possible origin for these sensory prediction signals is frontal cortical areas, which are involved in the initiation or control of vocal production. The role of the frontal cortex in human speech has been long understood, with areas exhibiting speech-related neural activity as well as causal roles in the control of speech timing and articulation.^31–38^ In contrast, the role of the frontal cortex in non-human primate vocalization has been controversial.^39–41^ However, activity in premotor and prefrontal areas, homologous to those in humans, have been found to exhibit neural activity before and during vocal production.^42–44^ Whether or not these frontal cortical areas provide vocal sensory prediction information or are involved in vocalization-induced suppression of the auditory cortex is unknown. A similar role of motor cortex in non-vocal behaviorally-induced suppression of the auditory cortex has been reported.^45–47^

Here, we sought to address these questions by simultaneously recording neural activity from both the auditory and frontal cortices of marmoset monkeys while they produced self-initiated vocalizations in the setting of natural communication. We found an increase in the theta-band power of the local field potential (LFP), as well as multi-unit neural activity, in both brain areas shortly before the onset of vocal production. During this pre-vocal period, there was an increase in directed signaling, measured with Granger causality, from the frontal cortex to the auditory cortex. This pre-vocal activity and signaling predicted both suppression of the auditory cortex as well as the acoustic properties of following vocalizations, suggesting the transmission of a predictive signal to the auditory cortex.

## Results

### Pre-vocal increases of theta-band power and multi-unit activities were seen in both auditory and frontal cortices

We recorded neural activity from the auditory (AC) and frontal cortices (FC) while monkeys freely produced vocalizations in their home colony (2560 AC recording sites and 1968 FC sites from three monkeys). AC recording sites included core auditory cortex (A1) as well as lateral- and para-belt regions. FC recording sites were estimated to be in the ventrolateral part of the frontal cortex, including Brodmann Areas (BA) 6, 8, 45, and 47.

Time-frequency analysis of local field potentials (LFPs) and multi-unit responses (MUA) in the AC revealed population-average suppression during vocal production, coupled with an increase in gamma-band LFP power (25-50 Hz), consistent with previous results^48^ (Fig. 1A, top). Frontal recordings showed a similar pattern of MUA and broadband LFP suppression during vocal production. Examination of the pre-vocal period, however, revealed an increase in low frequency theta-band power (4-8 Hz) preceding vocal onset by 0.5 to 1 seconds, seen in both AC and FC.

**Figure 1.**
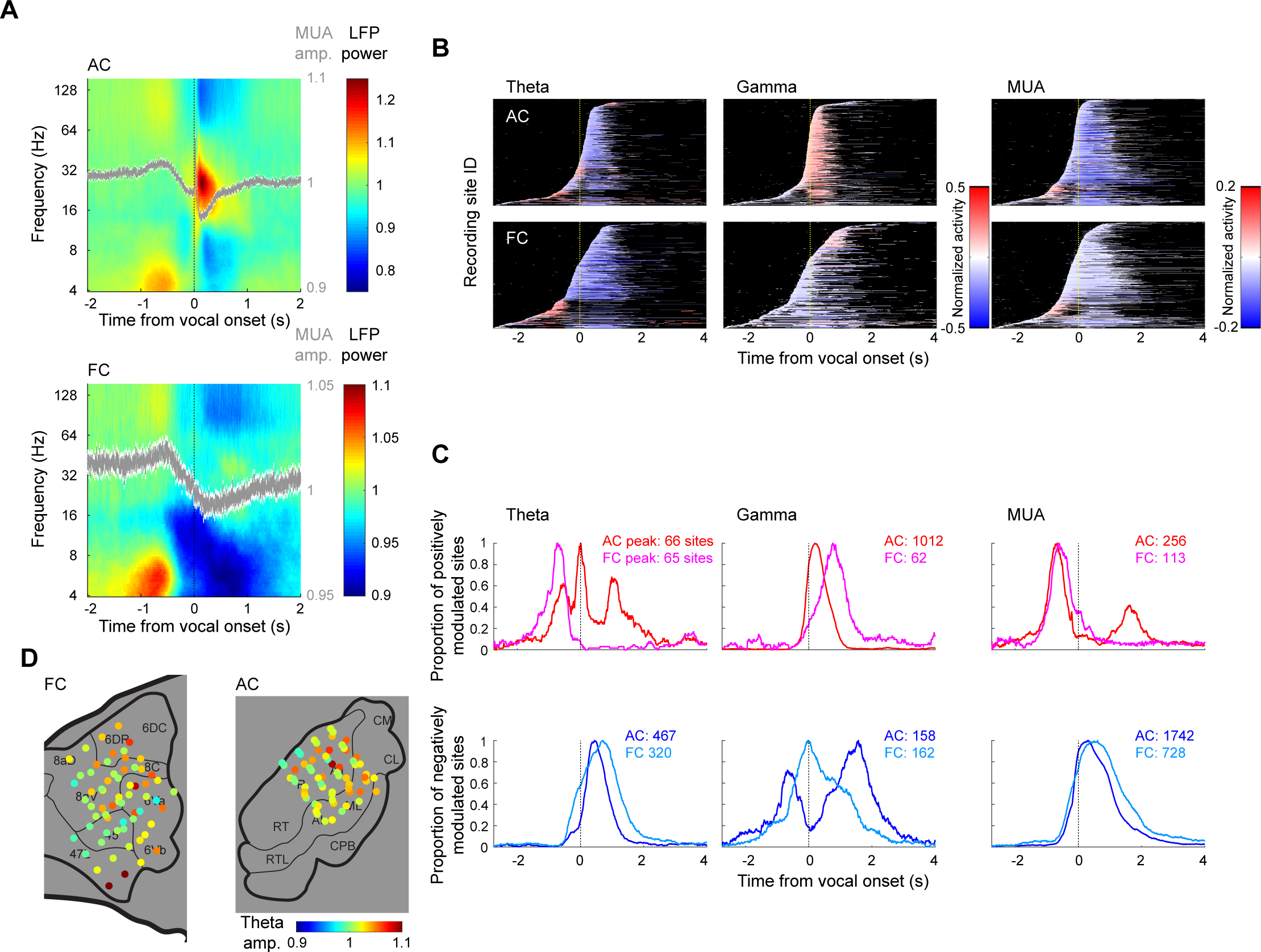
Vocalization-induced activity in frontal and auditory cortices. (**A**) Time-frequency plots of global average local field potential (LFPs) and multi-unit activity (MUA) recorded from all auditory (AC) and frontal cortex (FC) sites: 2,560 AC sites (top, 864 sites from monkey A, 832 from monkey Z, and 864 from monkey C) and 1,968 FC sites (bottom, 624 from monkey A, 480 from monkey Z, and 864 from monkey C). Color indicates LFP time-frequency power normalized by the median baseline power. Normalized mean MUA responses are overlaid (gray lines; Std error: white). (**B**) Time histogram of significantly modulated activity for theta (left) and gamma-band amplitude (middle), and MUA responses (right) are shown separately for AC (top) and FC (bottom). Data from individual recording sites are sorted by the onset time of modulated responses (Wilcoxon signed-rank test, p<0.05, FDR corrected). Color indicates the degree of the change compared to baseline amplitude (red: increase, blue: decrease). (**C**) Normalized proportions of significantly modulated sites are shown relative to vocal onset time. (top: positively modulated sites, bottom: negatively modulated sites). Numbers of sites with positive and negative modulations are indicated for both AC and FC (inset). (**D**) Anatomic distributions of all electrode recording sites colored by average pre-vocal (from 1 to 0.5s before vocalization) theta-band amplitude in FC (left) and AC (right). Data from both hemispheres are projected over the left hemisphere.

To better understand the timing of vocalization-related activity, we measured significant changes in theta, gamma, and MUA responses for individual recording sites In both AC and FC, we found that about 25% of recording sites showed changes in theta-band activity before and/or during vocalization (AC: 676/2560 [26%]; FC: 472/1968 [24%]). Importantly, a subset of these changes appeared in the pre-vocal period (AC: 285/2560 [11%] or 285/676 [42%] of modulated sites]; FC: 289/1968 [15%] or 289/472 [61%]), and their positive changes contributed to pre-vocal activity as seen in the population time-frequency analysis (Fig. 1A). We also found a large proportion of sites with pre-vocal increases that did not show significant changes during vocalization (AC: 65/126 [52%]; FC: 22/91 [24%]), suggesting the involvement of different neural populations in pre-vocal and vocal processing.

We also quantified site-by-site activity modulations in gamma-band and multi-unit activity (MUA). For gamma-band activity, consistent with our previous work^48^, half of recording sites in AC showed significant changes (1290/2560 [50%]), mostly with increased power (997/2560 [39%] or 997/1290 [77%] of modulated sites), while the proportion of gamma-band responses in FC was lower (315/1968 [16%]) and more likely to be negative (232/1968 [12%] or 232/315 [74%]). Also consistent with previous findings^17^, spiking activity measured by MUA was suppressed immediately before (∼200 ms) and during vocalization in AC (1559/2560 [61%] or 1559/1917 [81%]; Fig. 1A). Similar suppression during vocal production was also observed in FC, including pre-vocal suppression (750/1968 [38%] or 750/929 [81%]). Interestingly, we also found a subset of recording sites in AC (14% [347/2560] or 18% [347/1917] of modulated sites) and FC (9% [179/1968] or 18% [171/929]) showing pre-vocal increases in MUA followed by vocal suppression (∼1-0.5 s before vocalization; Fig. 1B), which was also visible in the population average responses (Fig. 1A). This pre-vocal increase was not as clearly evident in our past work, and aligned with the timing of the pre-vocal increase in theta-band activity.

To gain insights about the possible information flow between FC and AC, we compared the timing relationship of modulated LFP and MUA activity across areas, and found significant differences in the onset times for both positive and negative responses (Fig. 1B and C). The pre-vocal increased in theta-band activity in FC preceded that in AC (FC: 985 [1115-870] ms vs. AC: 235 [550-180] ms; median and 95% confidence intervals, Wilcoxon rank-sum test, p<0.001). Pre-vocal MUA increases were similar between FC and AC, though slightly earlier in AC (AC: 940 [980-920] ms; FC: 840 [870-790] ms, p<0.001). Theta and MUA increases during vocalization were less common, and almost exclusively seen in AC, and not FC, as were AC gamma-band responses (Fig. 1B and C, middle and right). Sites with suppressed activity showed generally similar timing patterns between AC and FC, and were mostly limited to suppression during, or immediately preceding, vocal production. It should be noted, however, that timing differences between different measures of activity may be difficult to directly compare because they reflect physiological processes with different temporal dynamics (e.g., low frequency theta-band oscillation is inherently slower than MUA). Nevertheless, the earlier modulation onset in FC theta-band could be consistent with FC providing inputs to AC before vocal production.

To better determine frontal sources of this vocalization-related activity, we examined the anatomic distribution of the observed responses, but did not find a systematic pattern in the cortical locations of sites with pre-vocal changes in frontal theta-band activity (Fig. 1D). However, AC theta responses showed an anterior-posterior (AP) dependency (linear regression: p<0.001) with greater pre-vocal theta responses in posterior sites, although no medial-lateral (ML) patterns were observed (p=0.92).

### The frontal cortex sends directed Granger signaling to the auditory cortex

To more directly test whether FC could provide inputs to AC before vocal production, we calculated Granger causality (GC) between pairs of recording sites, a technique which measures explainable variance of one neural response based upon another.^49,50^ We performed this connectivity analysis across multiple LFP frequencies and for different peri-vocal time periods (baseline, pre-vocal, and vocal), and found GC signaling primarily in the pre-vocal period, and with a bias towards FC to AC communication. Figure 2A shows an example pair of FC and AC sites exhibiting typical LFP responses, and GC_FC→AC_ (FC to AC) signaling increases in lower frequency bands during the pre-vocal period, but not during vocalization, compared to baseline connectivity. Interestingly, this same pair of sites did not exhibit GC_AC→FC_ signaling (AC back to FC). Population averages of multiple sites across all animals showed a similar pattern with pre-vocal increases in GC_FC→AC_, primarily in low frequencies such as the theta band, with little effects, or even decreases during vocalization (Fig. 2B,C; pre-vocal ΔGC in AC: mean ± SE: 5.2*10^−4^ ± 1.4*10^−4^, Wilcoxon sign-rank test, p<0.001; pre-vocal in FC, 3.2*10^−4^ ± 0.9*10^−4^, p<0.001; vocal in AC, −2.3*10^−4^ ± 0.5*10^−4^, p< 0.001; vocal in FC, −2.7*10^−4^ ± 0.7*10^−4^, p<0.001). Higher frequency bands, including alpha and beta bands, also showed significant but smaller increases in GC (Fig. 2A and B). Because the theta band exhibited the greatest increase in GC, coupled with an increased pre-vocal power, we focused primarily on GC in this band for further analysis.

**Figure 2.**
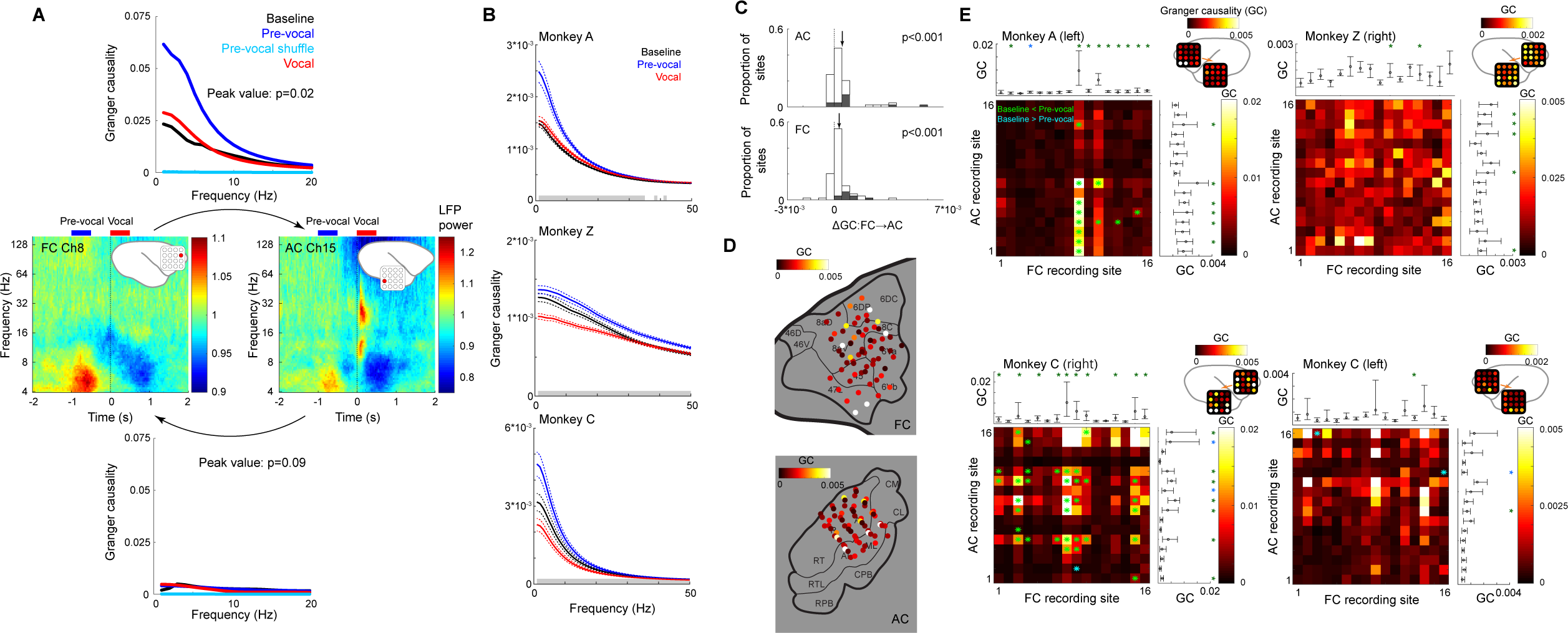
Directed Granger signaling from FC to AC in the pre-vocal period suggests top-down communication before vocal production onset. (**A**) Time-frequency plots are shown for a sample pair of AC and FC recording sites. Frequency-dependent, directional Granger causality (GC) values were calculated between these sites for both GC_FC→AC_ (top) and GC_AC→FC_ (bottom). Compared to baseline, there was an increase in the pre-vocal GC_FC→AC_ (−1 to −0.5 sec, blue), but not during the vocalization itself (red). Shuffled pre-vocal GC, controlling for general power increases, did not show similar results (light blue). Peak GC values were significantly different between time periods (Friedman test, p=0.02). Similar changes in GC_AC→FC_ (bottom) were not as apparent as GC_FC→AC._ (**B**) Population average GC_FC→AC_ values are shown across LFP frequencies, calculated separately from each animal, demonstrating increases in low frequency GC during the pre-vocal but not vocal period. Median and 95% confidence intervals are shown. Grey lines: frequency bins with significant differences between different time periods (p<0.05, Friedman test with FDR correction). (**C**) Population distributions of GC_FC→AC_ changes (ΔGC) during the pre-vocal period from baseline GC for sites in AC (top) and FC (bottom). Shaded bars indicate sites with significant changes (Wilcoxon signed-rank test, p<0.05). Arrows indicate mean values. (**D**) Spatial distribution of average pre-vocal GCs for recording sites in both FC and AC, showing greater values in a few area 47 and 6 sites of FC but no systematic pattern in AC. (**E**) Plots showing pre-vocal GC values for combinations of AC and FC sites for each animal and hemisphere. Averaged GC values and error bars (95% confidence interval) for each electrode track are shown in top (FC) and right (AC) marginal plots and on the cortical surface (right top). GC differences from baseline were statistically tested with Wilcoxon signed-rank test (p<0.05, FDR corrected) and reported with asterisks (green: pre-vocal increase; blue: pre-vocal decrease). GC values across AC (FC) sites were also compared using the Friedman test (p<0.001 for all monkeys), demonstrating heterogeneity of directed signaling between sites.

Although we identified population average increases in GC_FC→AC_ connectivity preceding vocalization, there was considerable heterogeneity between individual sites in both FC and AC (Fig. 2C). Examining the anatomic location of recording sites did not reveal any systematic distribution patterns in FC, although some of the strongest GC sites were located in ventral area 47, as well as more dorsal area 6 (Fig. 2D). Distribution patterns in AC also did not show systematic AP and ML dependencies (linear regression for AP: p=0.20; for ML: p=0.39). We examined whether specific combinations of FC and AC sites showed different degrees of GC_FC→AC_ connectivity and found a pattern in which a few FC sites provided the strongest inputs to a number of sites in AC (Fig. 2E). For example, in monkey A, FC recording site 9 (Fig. 2E top left), which is located in BA 47, and FC site 8 in animal C, provided broad inputs to many AC sites (Fig. 2E bottom left). These findings suggest that a specific subset of FC locations or areas are likely responsible for vocal FC to AC communication, but that they project broadly throughout the AC.

### Granger signaling and vocal suppression in auditory cortex

What information does this pre-vocal top-down GC_FC→AC_ convey? Given the vocal suppression seen in auditory cortex, one possibility is that this FC to AC communication reflects sensory prediction signals related to vocal production. To test this hypothesis, we examined the relationship between pre-vocal GC_FC→AC_ and spiking suppression in AC, which may encode sensory prediction and error signals during vocal production.^11,21^ We found a significant correlation between the pre-vocal GC_FC→AC_ signaling and the subsequent suppression of MUA responses, with AC sites receiving the largest GC being those with greatest decreases in MUA (Fig. 3). Interestingly, this was most evident in the pre-vocal period and immediately before vocal production, a time in which pre-vocal suppression begins (Fig. 3A middle and 3B; Spearman’s rank correlation, r=-0.09, and r=-0.12, p<0.001). We did not find a significant correlation between sites’ GC and the MUA during the vocalization itself (Fig. 3A, right), though this correlation returned around the time of vocal offset (Fig. 3B, right). This difference between pre-vocal and vocal MUA correlations may be a result of the integration of both vocalization-induced suppression and activity resulting from sensory inputs during vocal production that has been seen in some AC neurons.^17–19^ Overall, however, these correlations between GC and spiking suppression support the possibly contribution of FC pre-vocal inputs to vocalization-induced suppression in AC.

**Figure 3.**
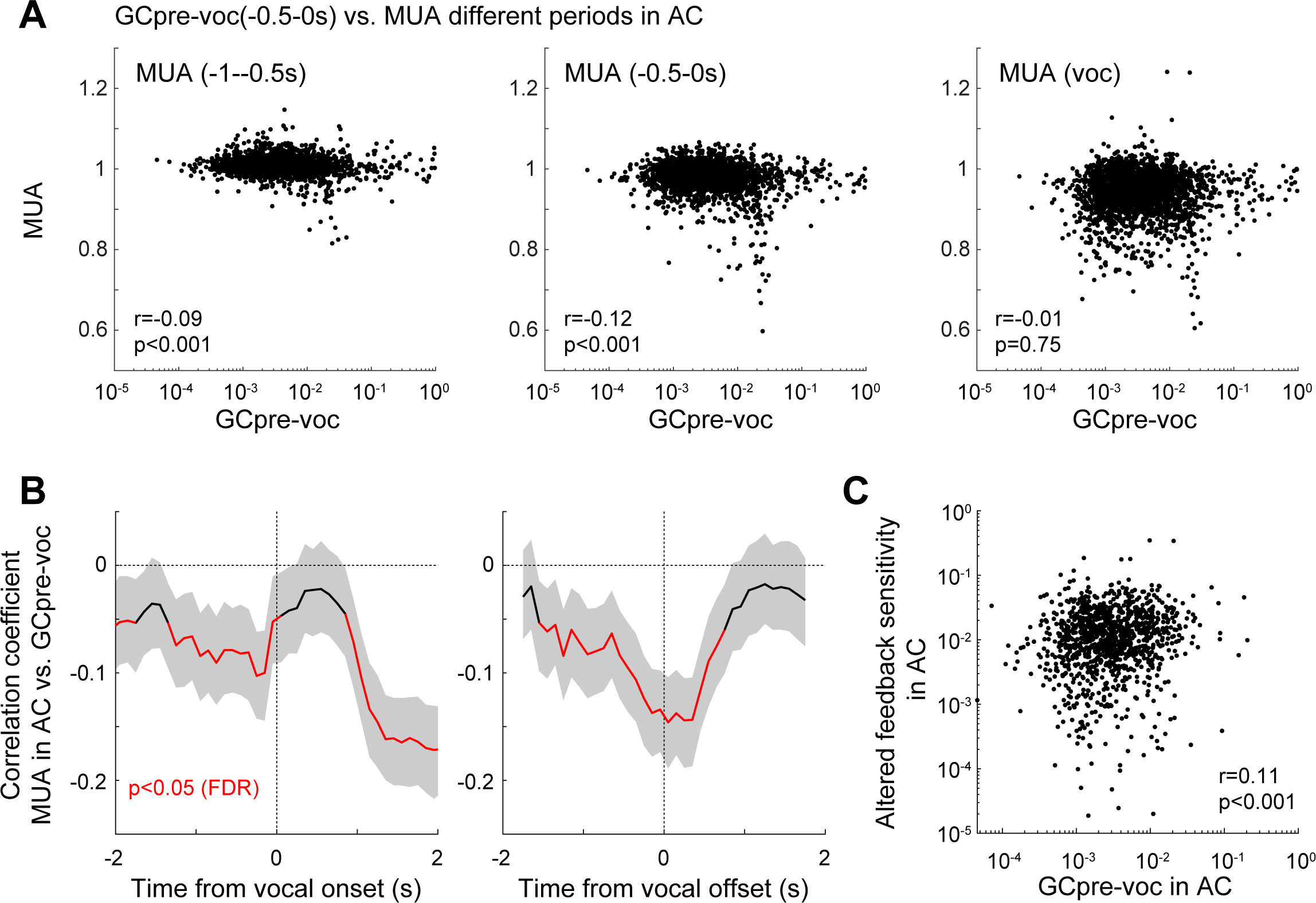
Granger causality predicts vocalization-induced suppression in the auditory cortex. (**A**) Scatter plots comparing pre-vocal GC_FC→AC_ in AC and multi-unit activity during different time periods: −1--0.5s (pre-vocal, left), −0.5-0s (middle), vocalization (right). Significant inverse correlations with MUA were seen during the early and immediate pre-vocal periods (p<0.05, Spearman rank correlation) but not for MUA during the actual vocal production period. (**B**) Time course of the GC-MUA correlation coefficients before and during vocal production, aligned by vocal onset (left) and offset (right). GC was fixed to the pre-vocal period, but compared to the time-varying MUA. Mean and 95% confidence intervals are plotted. Red lines: time bins with significant GC-MUA correlation (Spearman, p<0.05, FDR corrected). (**C**) Scatter plot showing a correlation between pre-vocal GC_FC→AC_ and sensitivity to altered (frequency-shifted) vocal feedback in AC sites during vocal production. Feedback sensitivity was calculated as the difference in MUA during vocalization between altered feedback and normal conditions. Spearman rank correlation is indicated.

Because of the putative role of vocal suppression in sensory prediction and vocal self-monitoring, we also examined whether AC sites with high GC_FC→AC_ connectivity were also more sensitivity to sensory prediction errors induced by altered vocal feedback. Comparing pre-vocal GC inputs to the differences between MUA responses during normal vocalizations and those with frequency-shifted vocal feedback, we found AC sites with greater GC_FC→AC_ inputs exhibited greater sensitivity to altered feedback (Fig. 3C; r=0.11, p<0.001). These results suggest AC sites receiving the strongest inputs from FC in the pre-vocal period were those that both were most likely to be suppressed and most likely to be sensitive to altered vocal feedback, consistent with a predictive signaling from FC for vocal production.

### Pre-vocal activity in both frontal and auditory cortices predicts vocal acoustics

Although the presence of top-down GC_FC→AC_ signaling, and correlation with vocal suppression, suggests that FC sends sensory prediction signals to AC, the nature of this signaling and information contained is still unclear. Motor prediction models for error detection suggest that such connections should convey specific predictions about the expected acoustics/sensory consequences of action.^29,30,47^ We therefore examined whether responses and connectivity in FC and AC encode information about the structure and acoustics of the vocalizations produced, focusing on vocal fundamental frequency (*f*0) as well as loudness (SPL) and duration.

Figure 4A shows an example FC recording site, comparing pre-vocal theta amplitude and the f0 of the following vocalization and exhibiting a significant correlation (r=-0.22, p=0.02). Across the population, we found that about 16% of FC sites showed a significant acoustic correlation between *f*0 and pre-vocal theta-band activity (mean correlation ± SE: 0.040 ± 0.002, p<0.001; Fig. 4B), and 11% for AC sites (0.012 ± 0.002, p<0.001). Looking at the timing of this correlation between neural activity and vocal f0, FC showed a large peak in both theta and MUA correlations in the pre-vocal period, maximal at the time of the pre-vocal theta band peak, with weaker correlations during the vocalization itself (Fig. 4C-E). Because this activity preceded vocal onset, it could not be a result of a sensory response, but rather likely reflects pre-motor or other signals predictive of the subsequent vocal production. In contrast to FC predictions, AC correlations showed two peaks, including a pre-vocal peak and a second peak during vocalization. This second AC correlation peak was often stronger than that pre-vocally (Fig. 4D), and present at more recording sites (Fig. 4E), which may reflect sensory responses to vocal sensory inputs. Interestingly, we observed significant correlations even before the pre-vocal period (Fig. 4C-E), implying the presence of background correlation between neural activity and acoustics, which could potentially reflect an underlying behavioral or emotional state. Nevertheless, the presence of correlation peak in the pre-vocal period that overlapped vocalization-specific neural activity suggests a more specific relationship between neural activity and vocal acoustics. We found similar pre-vocal correlations between theta-band activity and other vocal acoustic parameters, including vocal duration and SPL (vocal duration: AC: −0.011 ± 0.002, p<0.001; FC: −0.008 ± 0.002, p<0.001; SPL: AC: 0.005 ± 0.002, p=0.03; FC: −0.020 ± 0.002, p<0.001).

**Figure 4.**
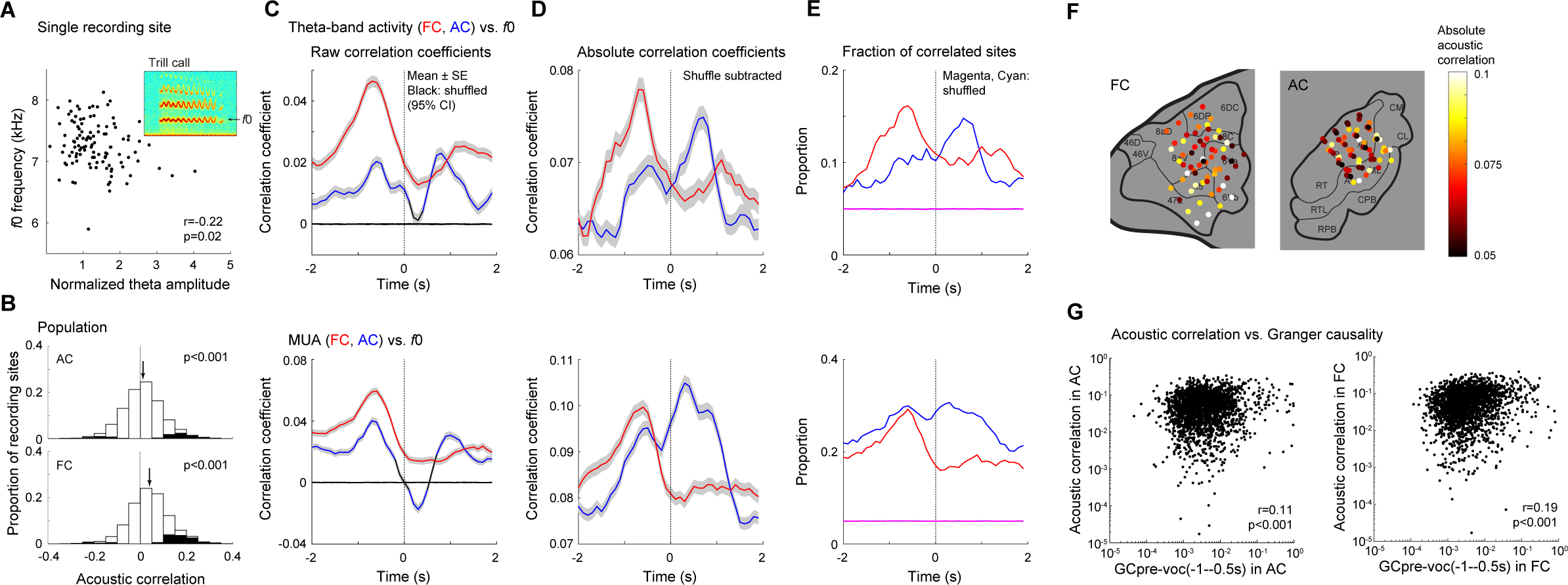
Relationship between neural activity, Granger causality, and vocal acoustic variability. (**A**) Scatter plot of a sample FC recording site demonstrating a correlation between pre-vocal theta-band activity and trial-to-trial fluctuations in vocal fundamental frequency (*f*0). (**B**) Population distributions of correlation coefficients between pre-vocal theta-band activity and *f*0 (acoustic correlation) for both AC and FC. Shaded bars indicate sites with significant correlations (p<0.05, Spearman). Arrows indicate mean values. (**C**) The time courses of population average acoustic correlations are shown for both FC (red) and AC (blue) sites, and separately for theta band-power (top) and MUA (bottom), demonstrating significant correlations before and during vocal production. Mean and standard error are plotted, as is a shuffle-corrected control (grey). Results are separately plotted as the mean absolute correlations (rectifying sites that inversely correlated with f0, **D**) and for the fraction of recording sites that individually exhibited significant correlations with vocal acoustics (**E**). (**F**) Anatomic distribution of acoustic correlations (absolute value) between pre-vocal theta-band power and vocal f0. (**G**) Scatter plot showing a correlation between pre-vocal GC_FC→AC_ and vocal acoustic correlation coefficients for sites in both in AC (r=0.11, p<0.001) and FC (r=0.19, p<0.001).

We further examined the anatomic distribution of pre-vocal theta band predictions/correlations with vocal acoustics (Fig. 4F). In FC, we noted a trend towards strong correlations in more ventral areas, though this may have also reflected stronger predictions from a specific animal. We did not find any pattern of correlation in AC, either for AP (linear regression, p=0.45) or ML (p=0.21) position. To evaluate whether this correlation of vocal acoustics could reflect a predictive top-down signal with specific acoustic information, we compared pre-vocal theta-acoustic correlations coefficients with GC_FC→AC_ at the same recording sites (Fig. 4G). Both AC and FC showed significant correlations between these measures (AC, r=0.11, p<0.001; FC, r=0.19, p<0.001), suggesting those sites sending (or receiving) top-down signals are also those whose activity most strongly varies with vocal acoustics, and therefore possibly reflecting the transmission of this sensory-motor prediction information.

### Directed signaling from AC to FC was decreased during vocal production

In contrast to the significantly increased GC_FC→AC_ communication in the pre-vocal period, we found a different pattern for bottom-up communication from AC to FC (GC_AC→FC_). In the pre-vocal period, this GC_AC→FC_ was unchanged from baseline levels (Fig 5A; for AC, mean ΔGC ± SE: 7.0*10^−5^ ± 6.9*10^−5^, p=0.29; for FC, 1.3*10^−4^ ± 0.7*10^−4^, p=0.07). During vocal production, however, we found decreases in GC_AC→FC_, most prominently in the theta-band frequencies. (Fig. 5A,B; for AC, mean ΔGC ± SE: −3.3*10^−4^ ± 0.8*10^−4^, p<0.001; for FC, −4.2*10^−4^ ± 0.8*10^−4^, p<0.001). As with the top-down GC, these bottom-up GC_AC→FC_ varied between different AC-FC site combinations (Fig. 5C), but even those sites with strong overall GC were reduced compared to their baseline levels. Cortical distribution patterns of GC_AC→FC_ in AC did not show systematic AP and ML dependencies during pre-vocal (linear regression for AP: p=0.27; for ML: p=0.46) and vocal periods (AP: p=0.41; ML: p=0.15). Interestingly, direct comparison of top-down (GC_FC→AC_) and bottom-up (GC_AC→FC_) signaling in FC correlated in both pre-vocal and vocal periods (for pre-vocal, r=0.31, p=0.01; for vocal, r=0.31, p=0.01) whereas AC did not show significant correlations (for pre-vocal, r=0.17, p=0.18; for vocal, r=0.24, p=0.05). These suggest common sites in FC receive inputs from AC and send outputs to AC, but AC sites receiving inputs from and sending outputs to FC are distinct. We did not find any consistent changes in vocal GC_AC→FC_ with frequency-shifted vocal feedback.

**Figure 5.**
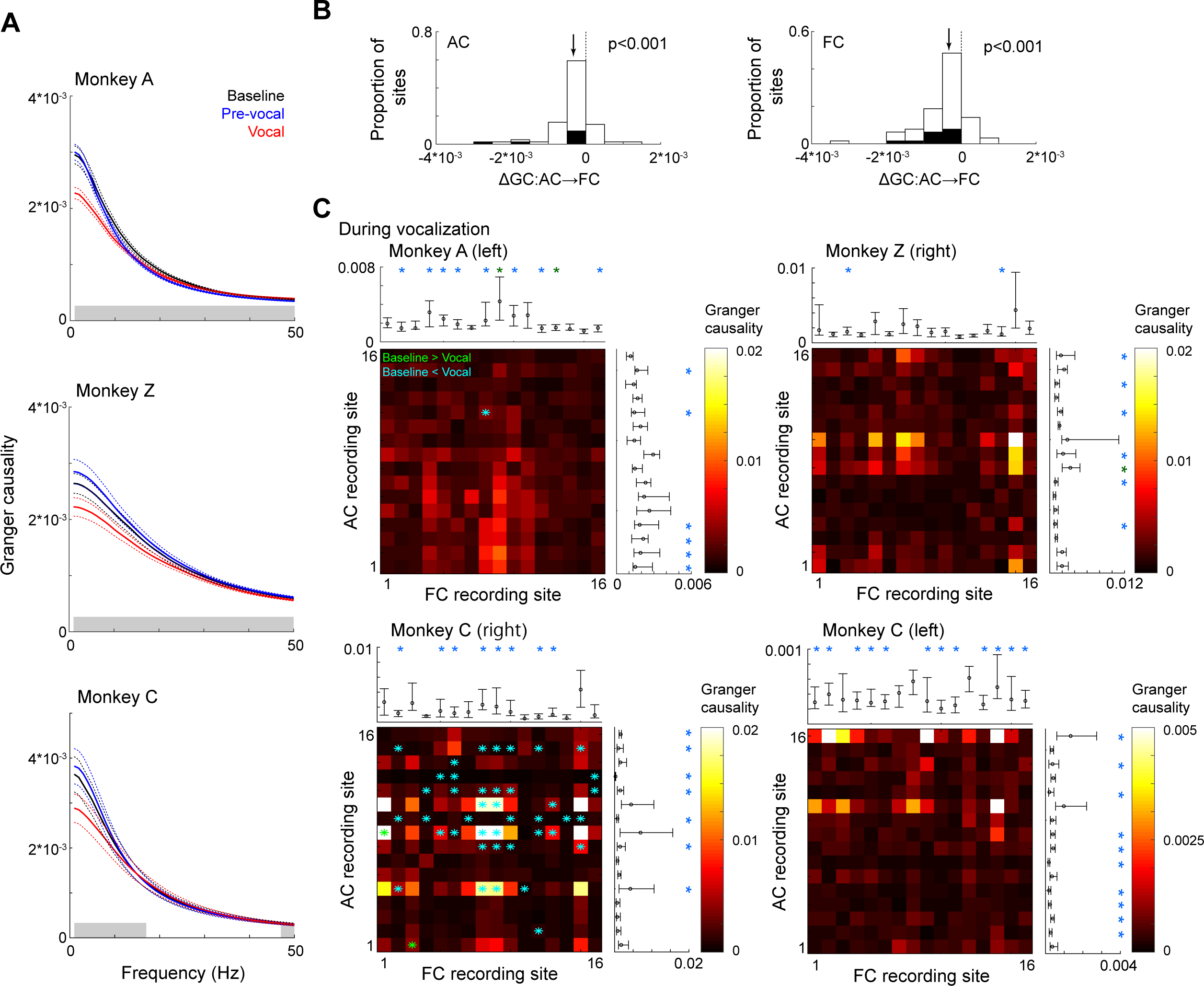
Weaker directed signaling from AC to FC during vocalization. (**A**) Average GC_AC→FC_ values across frequencies are compared between baseline, pre-vocal, and vocal time period. While there was decreased GC during vocal production, unlike GC_FC→AC_ there were no increases in pre-vocal GC. Grey lines: p<0.05 (Friedman with FDR correction). (**B**) Population distributions of GC_AC→FC_ changes (ΔGC) during vocalization from baseline GC in AC (left) and FC (right). Black bars indicate sites with significant changes (Wilcoxon signed-rank test, p<0.05). Arrows indicate mean values. (**C**) Plots showing vocal GC values for combinations of AC and FC sites for each animal and hemisphere. Averaged GC values and 95% confidence intervals for each recording site are shown in top (FC) and right (AC) marginal plots. GC differences from baseline were statistically tested with Wilcoxon signed-rank test (p<0.05, FDR corrected Wilcoxon) and reported with asterisks (green: increase; blue: decrease).

### Theta-band activity and directed signaling during vocal production are distinct from sensory processing

To test whether changes in directed signaling between FC and AC were specific for vocal production and not vocal sensory processing, we also calculated GC values during passively listening to playback of conspecific vocalizations (vocal playback; Fig. 6). In contrast to vocal production, time-frequency plots showed AC increased theta-band activity during vocal playback, but not in FC (Fig. 6A). Overall playback GC_FC→AC_ was smaller than GC calculated in the baseline period (Fig 6B top; mean ΔGC ± SE: −0.009 ± 0.003; p=0.03). This demonstrated a distinct pattern during playback compared to pre-vocal GC_FC→AC,_ which showed positive ΔGC in Fig 2C, although playback and pre-vocal GC_FC→AC_ correlated (r=0.09, p=0.001). In contrast, playback GC_AC→FC_ was similar to baseline GC (Fig 6B bottom, 0.002 ± 0.003, p=0.19), which is also different from the pattern seen during vocal production (i.e., Fig. 5B), suggesting the decrease of vocal GC_AC→FC_ is vocal-production specific, in contrast to the expected bottom-up communication seen during playback. For both GC_FC→AC_ and GC_AC→FC_, ΔGC measures were usually greater for playback than for pre-vocal (p<0.001), possibly due to differences in the recording environment (lower acoustic and electrical noises in the soundproof booth compared to colony) and behavior^51^ (head-restrained vs. free-moving).

**Figure 6.**
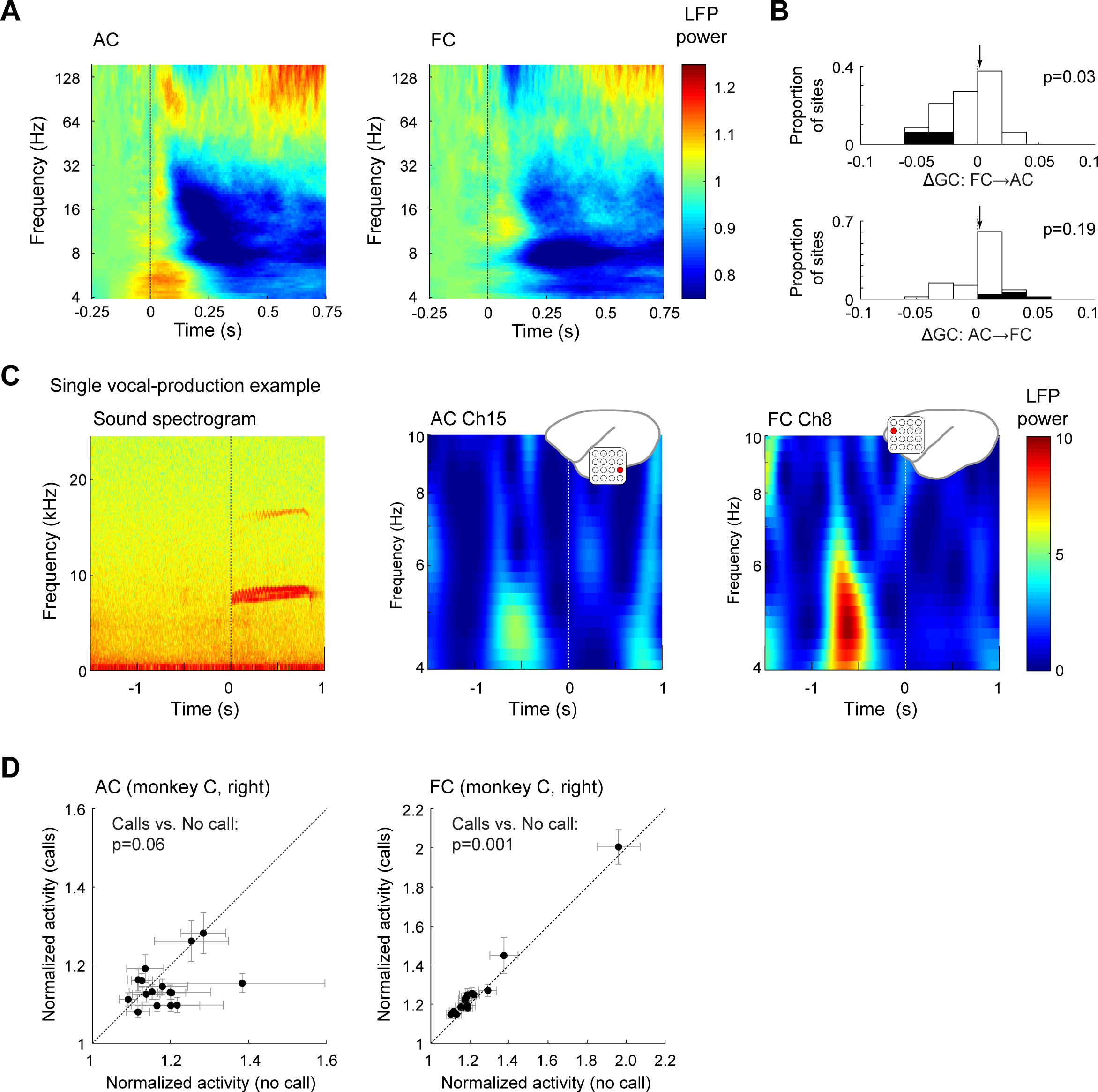
Sensory contributions to directed signaling and pre-vocal theta-band power. **(A)** Population average time-frequency plots of LFP power during passive listening to vocal playback (left: AC, right: FC), showing phasic increases of the theta-band and high frequency power in AC and broad band suppression in FC. These responses were distinct from the LFP responses seen during vocal production. (**B**) Distributions of GC changes (ΔGC) during vocal playback in AC (top) and FC (bottom). Shaded bars indicate significant changes (Wilcoxon, p<0.05). (**C-D**) Comparisons of LFP responses during spontaneous and interactive vocalizations. (**C**) Example sound spectrogram (left) and corresponding pre-vocal theta-band activity in AC (middle) and FC (right) for a sample recording site. Theta-band power increases and were noted even when there was no preceding vocalization. (**D**) Comparison of theta-band activity with vs. without preceding calls in AC (left) and FC (right) for a subset of sites. Site mean and standard error are indicated. Overall, there were similar pre-vocal responses whether or not there was a preceding vocalization, however there were small population-level increases in the presence of a preceding call (p=0.06 for AC and p=0.001 for FC, Wilcoxon).

Finally, to exclude the possibility that activity observed in the pre-vocal period was not simply a sensory response evoked by other monkeys’ calls, we analyzed a subset of our recording sessions, distinguishing vocalizations made in response to another vocalization from those with no preceding call in the pre-vocal period. We found similar increases in theta-band activity even in the absence of a preceding vocalization (Fig. 6C,D). Interestingly, while we did not find significant differences in pre-vocal activity between vocalizations with and without preceding calls in AC (with calls: 1.15 ± 0.01; without calls: 1.19 ± 0.02, Wilcoxon signed-rank test, p=0.06), we found a slight increase in FC theta with preceding calls (with calls: 1.27 ± 0.05; without calls: 1.24 ± 0.05, p=0.001). These results suggest that the pre-vocal activity cannot simply be explained by auditory response to others’ vocalizations.

## Discussion

In this study, we examined the presence of directed communication between frontal and temporal cortices during vocal production in order to determine if frontal cortex is a potential source for vocal sensory prediction signals previously seen in the auditory cortex. We found an increase in theta-band and muti-unit activity in both brain areas preceding the onset of vocal production by an average of 500 ms. This pre-vocal activity overlapped with an increase in top-down directional signaling from FC to AC and predicted the acoustics of subsequent vocalizations. More importantly, AC sites receiving FC inputs exhibited stronger vocalization-induced suppression, better acoustic predictions, and greater sensitivity to altered vocal feedback. These results suggest that FC may be the source of sensory prediction signals to the AC, beginning in the pre-vocal planning period. This interaction between FC and AC may be an underlying mechanism to calculate the error between intended and actual vocal outputs during vocal communication.

### Pre-vocal increases in the theta-band activity and MUA

In this study, we found a pre-vocal increase in both spiking (MUA) and LFP activity (theta-band) beginning 0.5-1 second before that start of vocal production. This was noted both in auditory as well as frontal cortex, though it had an earlier onset in frontal. In our previous work, we had described pre-vocal suppression in AC spiking activity beginning from up to a second before vocal onset, most notable in the last 0.2 s before vocal production.^11,17,19^ The presence of a pre-vocal peak in activity was not obvious in our previous studies, though there may have been a subtle hint of this finding (e.g., Fig 4 in Eliades and Wang, 2003^17^; Fig. 1 B and D in Tsunada and Eliades, 2020^48^). Several differences may explain why this pre-vocal increase was not emphasized previously. First was a difference in analytic time window; our previous studies often used 1-0.5 s before vocal production in the baseline period, while the current study defined the baseline period as 3-2.5 s before vocalization. Second, compared to pre-vocal suppression, the pre-vocal increase was more variable between recording sites, and only observed in a subset (Fig. 1 B and C; 80% for pre-vocal suppression and 10% for pre-vocal increase in MUA), and may have been masked in population level analysis. Finally, here we focused on multi-unit spike and LFP responses, rather than single unit, and previous reliance of single-units may have biased neural sampling.

In contrast to the AC findings, pre-vocal increases in MUA and spiking have been previously reported in marmoset FC^42–44^, although the timing of activity modulation was slightly earlier in our study (0.5-1 second here, up to 0.5 seconds in others’). The timing difference may have been a result of recording site differences, as our frontal sites were more anterior and lateral than in other studies, although we did not observe a systematic variation in the pre-vocal increase, or its timing, between frontal sites. Future studies will be needed to test functional differences in pre-vocal activity among FC sites, directly comparing rostral and caudal areas, as well as the use of perturbation techniques.

Similar pre-vocal increases in FC activity have also been observed in humans during word repetition. Specifically, Broca’s area, which presumably overlap our more ventral frontal sites, shows increases in the broadband LFPs (from ∼20 Hz to 150 Hz) ∼0.5 s before speech production^35^, although the primate analogue of Broca’s remains uncertain.^41,52,53^ However, because of the task design (repetition) in human studies, it is unclear if this increase represented a sensory response vs. preparation for speech production. Our finding of the pre-vocal activation in AC and FC, even in the absence of-preceding vocalization sound inputs, demonstrated that this activity is not simply a sensory response to auditory inputs. Instead, the correlation between the pre-vocal increase in theta-band activity and acoustics of following vocalization suggests that pre-vocal activity in AC and FC may encode sensory prediction signal of producing vocalization.

### Frontal cortex conveys sensory prediction information to auditory cortex

Our primary finding in this study was the presence of top-down communication from FC to AC during the pre-vocal period. This directed signaling overlapped with the pre-vocal increase in LFP theta-band activity, common to both AC and FC, and was strongest in the same low-frequency bands. Interestingly, it appeared that a subset of FC sites provided much of this input to a much broader set of sites in AC, including both the primary and higher order auditory cortex. There was no specific pattern to the distribution of the frontal sites, although the sites with the strongest were generally in the more ventral areas (putative Broca’s homolog), and in some of the more posterior-dorsal areas (possibly pre-motor cortex). The presence of connectivity to broad areas of AC was, perhaps, unexpected, given that reciprocal frontal-temporal connections have generally been described with belt rather than core regions of auditory cortex.^54–59^ One possible explanation is that temporal dynamics of the theta-band activity are relatively slow, and therefore theta-band interactions may be observed from both direct and indirect interactions between brain areas.^60,61^

Human studies have similarly revealed functional connectivity in LFP coherence between AC and FC during speech production compared to speech perception.^33,62^ In contrast, more recent studies using intracranial recordings have found information flow from Broca’s area to AC observed before but not during speech production.^35,63^ Similar pre-vocal interaction has also been found in the AC-FC circuits of bats.^64^ These findings suggest that Broca’s area interacts with AC specifically in the pre-vocal period, whereas other frontal areas may interact with AC even during vocal production. Although we do not know to what extent our recording sites are in areas homologous to Broca’s area, though possibly including our more ventral electrodes, our results for pre-vocal interaction between AC and FC are consistent with the pre-vocal interaction between Broca’s area and AC.

What sort of information could this frontal-auditory interaction convey? Although Broca’s area has been suggested to play a role in articulatory programming and controlling speech timing^35,36,63^, it is unclear what role activity in marmoset frontal activity plays during vocal production and, therefore, what sort of top-down information might be conveyed. The role of frontal cortex in non-human mammalian vocalization has long been controversial, and it is unclear to what degree it is involved in specific motor control of vocal production.^39,40,65^ Our call-by-call analysis revealed that pre-vocal activity was actually predictive of subsequent vocalization acoustics, suggesting the representation of a sensory prediction signal in FC (and AC). An alternative explanation would be that this activity reflected some sort of motivational internal state or arousal, which itself was then correlated with an animal’s vocal behavior. The time-locked pre-vocal increase in this acoustic correlation would seem to argue against a more general internal state, but is difficult to ambiguate without more direct measures of arousal. Additionally, we also found a relationship between the top-down directional signaling and altered feedback sensitivity in AC, which likely reflects sensory prediction error. Together, these findings are consistent with a model in which FC conveys preparatory and sensory prediction signals to AC before vocal production and then, during vocal production, AC sites receiving this input play a role in calculating sensory error between intended vocalizations and actual vocal outputs.

### Mechanism to control ongoing vocal production

Although we found greater FC-AC interactions in the pre-vocal period, we found weaker interaction during vocal production itself, and little signaling from AC back to FC. These findings may be puzzling given monkeys’ ability to control ongoing vocal production, a behavior that likely depends on top-down sensory prediction during vocal production^11^, and may also involve reciprocal projections of error information from AC back to the vocal motor system. One possible explanation is that our FC sites were fairly anterior, and it is possible that intermediary stages of processing are involved in ongoing frontal-temporal interaction once vocalization actually begins. Putative areas could include motor, pre-motor, and supplementary motor areas that have been described^31,32,34,36,44,66^, but were likely not covered by our electrode arrays. The more anterior and ventral areas we sampled may have been more involved in pre-motor planning, and then relayed information to motor areas which were then responsible for ongoing control and sensory prediction during the actual production, as well as receiving the error signal outputs of auditory cortex. Similar motor to sensory projections have been found to be responsible for locomotion-induced suppression in mice.^45–47^ Human ECoG and cooling studies have also supported such a model, with different frontal cortical areas responsible for speech timing, planning, and ongoing articulation control.^36,37^ It is also possible that vocal error outputs project to brain areas beyond cortex, including cerebellum, basal ganglia and brainstem nuclei involved in vocal motor control.^39,67^ Future studies will need test the underlying circuit mechanisms for sensory prediction and error signal transmission between AC and frontal motor areas, as well as these subcortical structures.

### A possible role of directional signaling from FC to AC in social vocal communication

More generally, the importance of the top-down signal from FC to AC has also been emphasized in predictive processing during speech perception.^68–70^ Briefly, theta oscillations in AC are entrained by the rhythmic component of speech (3-8 Hz). This coupling can be further facilitated by top-down signals from FC, possibly through temporal gating and content expectation. Interestingly, marmosets exhibit a similar theta rhythm in their phonatory and articulatory systems during vocalizations, suggesting the common theta-band entrainment during vocal perception.^71^ Together with our findings of theta-band interactions between FC and AC for vocal production, vocal perception and production may share some common theta-band communication mechanisms, including efficient neural communication by entraining both brain areas to quasi-rhythmic components of speech.^69,72–75^ Although we did not find similar FC-AC interactions during passive listening conditions, it would be interesting to compare neural interactions in the same communicative context, where perception is a more active process, or during behavioral tasks, given the neural activity highly depends on the behavioral context.^51,76^

## Materials and Methods

Experiments were performed using three adult marmoset monkeys (*Callithrix jacchus*). One animal had also been used in previous studies of the auditory cortex.^11,48^ We simultaneously recorded neural activity in both auditory and frontal cortices using implanted multi-electrode arrays during natural, self-initiated vocal production. All experiments were conducted under the supervision and approval of the University of Pennsylvania Animal Care and Use Committee.

### Vocal and neural recording

As in our past work, vocal recordings were performed with the subject animal in a small cage with a custom three-walled sound attenuation booth to improve sound quality, which allowed to visually and vocal with other animals in the colony.^19,21,77^ We recorded vocalizations using a directional microphone (Sennheiser ME66) placed ∼20cm in front of the monkey using an amplifier (Focusrite OctoPre MkII) and data storage system with 48.8 kHz sampling rate (RX-8, Tucker-Davis Technologies). Vocalizations produced by other monkeys in the colony were captured using additional microphones (C1000S, AKG). Offline analysis extracted vocalizations from the recorded signals and classified them into established marmoset call types based on their spectrograms using a semi-automated system^78^ (see the details in Vocal acoustic analysis). Although all major call types were produced by the subject animals, we analyzed phee, trillphee, and trill calls because these calls were the most commonly produced and previously found to have similar vocal suppression within individual units.^19^

We recorded neural activity with implanted multi-electrode arrays (Warp 16, Neuralynx) in the auditory and frontal cortices (Monkey A: right auditory cortex and left auditory and frontal cortices; Monkey Z: right auditory and frontal cortices and left auditory cortex; Monkey C: right and left auditory and frontal cortices; Eliades and Wang, 2008). The arrays consist of a 4×4 grid of individually moveable electrodes (4 Mohm tungsten, FHC). Neural signals were sampled at 24 kHz and stored for offline analysis (RA16CH, RZ2, and RS4, Tucker-Davis Technologies). Here, we primarily focused on the analysis of population activity, including multi-unit activity (MUA) and local field potentials (LFPs). We extracted MUA and LFPs by first subtracting the average activity recorded from all electrodes on an array to reduce muscle potentials and other movement artifacts, and then band-pass filtering (300-5000 Hz for MUA and 1-300 Hz for LFPs) and down-sampling to 1 kHz. We chose and LFP-based analysis because they may be more suitable to detect interactions across cortical areas and have been previously used for these sort of long-range connectivity analyses.^79–82^ More practically, because single-unit activity is significantly suppressed during vocalization in AC, statistical analysis of spike-based connectivity would be more challenging due to the limited number of data points.

Prior to each experimental session, we also characterized responses to passive playback of vocal sounds. Stimuli were delivered through a speaker (B&W 686S2) located 1m in front of a chair-restrained animal within a sound booth (Industrial Acoustics, Bronx NY). We presented multiple vocalization stimuli, including samples of animals’ own vocalizations (previously recorded from each animal) and conspecific vocalization samples (from other unknown individuals) at loudness levels matching those produced in the colony recording.

### Altered auditory feedback

For a subset of neural recording sessions, we manipulated auditory feedback of produced vocalizations, similar to our past work.^11^ Real-time manipulation was conducted by modifying the vocal signal to shift the frequency by ±2 semitone through a commercial effects processor (Eventide Eclipse V4). The sound intensity of the shifted feedback was calibrated (Crown XLS1000) to ∼10 dB sound pressure level (SPL) above the intensity of direct, air-conducted feedback. Altered feedback signals were returned to the animal through earbud-style headphones (Sony MDR-EX10LP) modified to attach to the animal’s head post (Eliades and Wang, 2008). Typically, we only shifted the feedback in one direction (either −2ST or +2ST) in any given recording session. During feedback sessions, we delivered shifted feedback only for a random subset of vocalizations (either 50 or 60% of vocalizations) to prevent adaptation or prediction. Technical details for vocal detection and triggering timing of shifted feedback have been discussed in our previous paper.^11^

### Vocal acoustic analysis

Vocalizations were first detected and extracted from recordings using a semi-automated method based on crossings of a sound amplitude threshold. Vocalizations were then classified into established marmoset call types based on visual inspection of the spectrograms.^78,83,84^ We next characterized the acoustic properties and call structure of each vocalization, including duration, fundamental frequency (*f*0), and vocal amplitude. For the *f*0, we first generated call spectrograms to identify a frequency with the maximum power in each time bin and then averaged the frequencies over time. The vocal amplitude was calculated as a root mean square amplitude averaged across the entire vocal duration.

### Anatomical localization of recording sites

Electrode array locations within the cortex were estimated using paraformaldehyde (4%) fixed brains following the completion of neural recordings and, in some cases, electrolytic lesions. Intact brain tissues were visually inspected, and photographs taken from several viewpoints, locating surface tissue effects from the arrays and lesions. Images were then scaled for size differences and aligned to the marmoset brain atlas (https://3dviewer.marmosetbrainmapping.org/). Array locations were overlayed with the atlas to estimate their corresponding Brodmann areas within frontal cortex.^43,44^

### Neural data analysis

**V**ocal production related LFPs were first visualized using time-frequency plots. We applied a Morlet wavelet transform (the width of Gaussian as 6, frequency range from 4 Hz to 150 Hz) for individual vocal response, followed by averaging across vocal events. Based upon known pre-vocal suppression^17^ and inspection of LFP and MUA time courses, we defined a baseline period as the time window spanning 3-3.5 s before vocal onset, a pre-vocal period of 1-0.5 s before, and a vocal period for the duration of each vocalization. LFP and MUA during pre-vocal and vocal time periods were normalized for each recording site by dividing by baseline activity. The time-frequency analysis of LFPs showed vocalization-related modulations in the theta- and gamma-band power. To quantify these changes, we extracted those bands’ amplitudes using Hilbert transformation after applying four-pole Butterworth filters (for theta band, 4-8 Hz; for gamma band, 25-50 Hz), and normalized them by baseline amplitudes. Similarly, we also extracted MUA amplitudes applying Hilbert transformation to the pre-processed MUA.

To determine the timing of peri-vocal changes in both MUA and specific LFP frequency bands at individual recording sites, we quantified responses using a sliding window of baseline-normalized activity:(R_bin_ – R_baseline_)/R_baseline_, where R_bin_ is the amplitude of LFPs or MUA for each time bin, and R_baseline_ is baseline activity. We used 500 ms time bins to match the baseline time window length, with shifting 10 ms step. The size of time bins (e.g., 200 ms) did not change results significantly. We determined the timing of significant responses using a running Wilcoxon rank-sum test with false discovery rate (FDR) correction to compare LFPs and MUA amplitudes at each time bin with baseline activity (p<0.05). We defined the onset of significantly modulated activity as the first of 10 consecutive time bins with reliably different responses from baseline activity.

Although we found modulated activity before vocal production (pre-vocal activity), we were concerned that this could reflect sensory responses to preceding vocalizations produced by other monkeys in the recording environment. To test the effect of the existence of preceding calls, we examined a subset of experimental sessions, first identifying vocalizations produced by other monkeys using audio recordings on a colony-facing microphone (away from the experimental subject). We classified each of a subject’s vocalization as with or without preceding calls in the pre-vocal period and compared pre-vocal activity between those two conditions. This analysis was conducted over 1790 calls obtained from 9 sessions.

For analysis of auditory feedback effects, we compared MUA responses for normal vs. shifted feedback conditions, calculating an absolute MUA difference during vocal production between those conditions. We pooled recording sessions with different directions of frequency shifts (i.e., −2ST and +2ST) based on our previous findings of similar population average responses for different shifts.^11^

To test the information flow and functional connectivity between AC and FC, we calculate Granger causality (GC) using the FieldTrip toolbox.^85^ Briefly, GC quantifies how the inclusion of past AC (or FC) signals improves the prediction of FC (or AC) signals compared to just using past FC signals.^49,80,86^ Importantly, GC values do not simply depend on the magnitude of the activity, but they reflect the predictability of a sequence of data from another. This was a deliberate choice due to concerns that the high degree of AC suppression during vocal production might otherwise affect the connectivity analysis. For the baseline, pre-vocal, and vocal production periods, we separately calculated GC for each frequency using event-induced activity after subtraction of event time-locked activity. We chose to calculate GC for each individual frequency instead of using the average LFP (across frequencies) because the average LFP is usually dominated by lower-frequency LFPs, and higher-frequency components and their interactions can be easily masked. In fact, band-specific interactions have been used in previous functional connectivity studies.^79–82^ We calculated GC values for all possible pairings of simultaneously recorded AC and FC sites during a given experimental session. We calculated an average GC value for each site, for each frequency (1-150 Hz), and for each electrode recording track, for further analysis and visualization of cortical distribution. A subset of analyses extracted GC values in the theta band. In order to test the statistical significance of GC values, we first compared GC values among the different time periods (baseline, pre-vocal, vocal) for each frequency using the Friedman test with FDR correction. Since this initial analysis showed greater pre-vocal GCs, we statistically compared pre-vocal vs. baseline GCs for each site and for each recording track (Wilcoxon sign rank test). As an additional statistical test, we also generated a control distribution for each AC-FC pair using call (or trial) shuffled data. Shuffled data was created 1000 times, and 95% confidence intervals were calculated. While most analyses used the raw GC result, some analyses also used the difference of GC values in the theta band between pre-vocal and vocal periods and baseline; we calculated ΔGC by subtracting baseline GC value from pre-vocal and vocal GC.

To test the correlational structures among vocalization-related behavior, neural activity, and functional connectivity patterns, we conducted several Spearman’s rank correlation analyses. To test the relationship between pre-vocal GC and time-varying MUA at the population level, we performed a running correlation analysis with FDR correction (500 ms time bins, 100 ms step). We also tested whether pre-vocal activity predicted acoustics (*f*0, call amplitude) and call structure (duration) of subsequent vocalizations. This acoustic correlation analysis was conducted for pre-vocal activity as well as time-varying for each site. At the population level, we examined the distribution of raw acoustic correlations values and the fraction of sites with significant correlation. In addition to raw correlation coefficients, we also calculated the absolute magnitude of coefficients given the possibility of similar functional roles for positive and negative vocal acoustic correlations. In general, we used p-values obtained from the rank correlation to measure statistical significance, but we also confirmed by generating a control distribution using shuffled data. For the acoustic correlation, we shuffled pairs of theta-band activity and acoustics (e.g., *f*0) and calculated correlation coefficients. Statistical significance was tested using 95% confidence intervals of the mean from 1000 times shuffling data.

To test the relationships between the anatomic location of recording sites and physiological measurements, including theta-band activity, GC values, acoustic correlation, we applied multivariate linear regression analyses as a function of animal, hemisphere, anterior-posterior position, and medial-lateral position.

Unless otherwise noted, all statistical tests were performed using non-parametric methods. P-values <0.05 were considered statistically significant.

## Acknowledgements

This work was supported by NIH grants DC014299 and DC018525 (S.J.E.), the Triological Society Clinician-Scientist Development Award (S.J.E.), the Ministry of Education, Culture, Sports, Science and Technology of Japan Leading Initiative for Excellent Young Researchers Grant 1071421 (J.T.), Grant A19K237690 (J.T.), Ichiro Kanehara Foundation (J.T.), and Daiichi Sankyo Foundation of Life Science (J.T.), and the National Natural Science Foundation of China (J.T.). We thank T. Coleman and P. Sayde for assistance in animal training and care.

## Author contributions

J.T. and S.J.E. designed research; J.T. and S.J.E. performed research; J.T. and S.J.E. analyzed data; J.T. and S.J.E. edited the paper; J.T. and S.J.E. wrote the paper.

## References

1. Houde, J.F., and Jordan, M.I. (1998). Sensorimotor adaptation in speech production. Science 279, 1213–1216.

2. Guenther, F.H., Ghosh, S.S., and Tourville, J.A. (2006). Neural modeling and imaging of the cortical interactions underlying syllable production. Brain Lang 96, 280–301. 10.1016/j.bandl.2005.06.001.

3. Hickok, G., Houde, J., and Rong, F. (2011). Sensorimotor integration in speech processing: computational basis and neural organization. Neuron 69, 407–422. 10.1016/j.neuron.2011.01.019.

4. Eliades, S.J., and Wang, X. (2019). Corollary Discharge Mechanisms During Vocal Production in Marmoset Monkeys. Biol Psychiatry Cogn Neurosci Neuroimaging 4, 805–812. 10.1016/j.bpsc.2019.06.008.

5. Lombard, E. (1911). Le signe de l’elevation de la voix. Annales des Maladies de l’Oreille et du Larynx 37, 101–119.

6. Lane, H., and Tranel, B. (1971). The Lombard sign and the role of hearing in speech. J Speech Hear Res 14, 677–709.

7. Brumm, H., and Zollinger, S.A. (2011). The evolution of the Lombard effect: 100 years of psychoacoustic research. Behaviour 148, 1173–1198.

8. Eliades, S.J., and Wang, X. (2012). Neural correlates of the lombard effect in primate auditory cortex. J Neurosci 32, 10737–10748. 10.1523/JNEUROSCI.3448-11.2012.

9. Luo, J., Hage, S.R., and Moss, C.F. (2018). The Lombard Effect: From Acoustics to Neural Mechanisms. Trends Neurosci 41, 938–949. 10.1016/j.tins.2018.07.011.

10. Burnett, T.A., Freedland, M.B., Larson, C.R., and Hain, T.C. (1998). Voice F0 responses to manipulations in pitch feedback. J Acoust Soc Am 103, 3153–3161. 10.1121/1.423073.

11. Eliades, S.J., and Tsunada, J. (2018). Auditory cortical activity drives feedback-dependent vocal control in marmosets. Nat Commun 9, 2540. 10.1038/s41467-018-04961-8.

12. Muller-Preuss, P., and Ploog, D. (1981). Inhibition of auditory cortical neurons during phonation. Brain Res 215, 61–76.

13. Creutzfeldt, O., Ojemann, G., and Lettich, E. (1989). Neuronal activity in the human lateral temporal lobe. II. Responses to the subjects own voice. Exp Brain Res 77, 476–489. 10.1007/BF00249601.

14. Paus, T., Perry, D.W., Zatorre, R.J., Worsley, K.J., and Evans, A.C. (1996). Modulation of cerebral blood flow in the human auditory cortex during speech: role of motor-to-sensory discharges. Eur J Neurosci 8, 2236–2246. 10.1111/j.1460-9568.1996.tb01187.x.

15. Numminen, J., Salmelin, R., and Hari, R. (1999). Subject’s own speech reduces reactivity of the human auditory cortex. Neurosci Lett 265, 119–122. 10.1016/s0304-3940(99)00218-9.

16. Houde, J.F., Nagarajan, S.S., Sekihara, K., and Merzenich, M.M. (2002). Modulation of the auditory cortex during speech: an MEG study. J Cogn Neurosci 14, 1125–1138. 10.1162/089892902760807140.

17. Eliades, S.J., and Wang, X. (2003). Sensory-motor interaction in the primate auditory cortex during self-initiated vocalizations. J Neurophysiol 89, 2194–2207. 10.1152/jn.00627.2002.

18. Eliades, S.J., and Wang, X. (2005). Dynamics of auditory-vocal interaction in monkey auditory cortex. Cereb Cortex 15, 1510–1523. 10.1093/cercor/bhi030.

19. Eliades, S.J., and Wang, X. (2013). Comparison of auditory-vocal interactions across multiple types of vocalizations in marmoset auditory cortex. J Neurophysiol 109, 1638–1657. 10.1152/jn.00698.2012.

20. Greenlee, J.D., Jackson, A.W., Chen, F., Larson, C.R., Oya, H., Kawasaki, H., Chen, H., and Howard, M.A., 3rd (2011). Human auditory cortical activation during self-vocalization. PLoS One 6, e14744. 10.1371/journal.pone.0014744.

21. Eliades, S.J., and Wang, X. (2008). Neural substrates of vocalization feedback monitoring in primate auditory cortex. Nature 453, 1102–1106. 10.1038/nature06910.

22. Greenlee, J.D., Behroozmand, R., Larson, C.R., Jackson, A.W., Chen, F., Hansen, D.R., Oya, H., Kawasaki, H., and Howard, M.A., 3rd (2013). Sensory-motor interactions for vocal pitch monitoring in non-primary human auditory cortex. PLoS One 8, e60783. 10.1371/journal.pone.0060783.

23. Behroozmand, R., Oya, H., Nourski, K.V., Kawasaki, H., Larson, C.R., Brugge, J.F., Howard, M.A., 3rd, and Greenlee, J.D. (2016). Neural Correlates of Vocal Production and Motor Control in Human Heschl’s Gyrus. J Neurosci 36, 2302–2315. 10.1523/JNEUROSCI.3305-14.2016.

24. Eliades, S.J., and Tsunada, J. (2023). Effects of Cortical Stimulation on Feedback-Dependent Vocal Control in Non-Human Primates. Laryngoscope 133 *Suppl 2*, S1–S10. 10.1002/lary.30175.

25. Sperry, R.W. (1950). Neural basis of the spontaneous optokinetic responses produced by visual inversion. J Comp Physiol Psych 43, 482–489.

26. Bridgeman, B. (1995). A review of the role of efference copy in sensory and oculomotor control systems. Ann Biomed Eng 23, 409–422. 10.1007/BF02584441.

27. Wolpert, D.M., Ghahramani, Z., and Jordan, M.I. (1995). An internal model for sensorimotor integration. Science 269, 1880–1882. 10.1126/science.7569931.

28. Poulet, J.F., and Hedwig, B. (2002). A corollary discharge maintains auditory sensitivity during sound production. Nature 418, 872–876. 10.1038/nature00919.

29. Crapse, T.B., and Sommer, M.A. (2008). Corollary discharge circuits in the primate brain. Curr Opin Neurobiol 18, 552–557. 10.1016/j.conb.2008.09.017.

30. Crapse, T.B., and Sommer, M.A. (2008). Corollary discharge across the animal kingdom. Nat Rev Neurosci 9, 587–600. 10.1038/nrn2457.

31. Bouchard, K.E., Mesgarani, N., Johnson, K., and Chang, E.F. (2013). Functional organization of human sensorimotor cortex for speech articulation. Nature 495, 327–332. 10.1038/nature11911.

32. Chang, E.F., Niziolek, C.A., Knight, R.T., Nagarajan, S.S., and Houde, J.F. (2013). Human cortical sensorimotor network underlying feedback control of vocal pitch. Proc Natl Acad Sci U S A 110, 2653–2658. 10.1073/pnas.1216827110.

33. Kingyon, J., Behroozmand, R., Kelley, R., Oya, H., Kawasaki, H., Narayanan, N.S., and Greenlee, J.D. (2015). High-gamma band fronto-temporal coherence as a measure of functional connectivity in speech motor control. Neuroscience 305, 15–25. 10.1016/j.neuroscience.2015.07.069.

34. Behroozmand, R., Shebek, R., Hansen, D.R., Oya, H., Robin, D.A., Howard, M.A., 3rd, and Greenlee, J.D. (2015). Sensory-motor networks involved in speech production and motor control: an fMRI study. Neuroimage 109, 418–428. 10.1016/j.neuroimage.2015.01.040.

35. Flinker, A., Korzeniewska, A., Shestyuk, A.Y., Franaszczuk, P.J., Dronkers, N.F., Knight, R.T., and Crone, N.E. (2015). Redefining the role of Broca’s area in speech. Proc Natl Acad Sci U S A 112, 2871–2875. 10.1073/pnas.1414491112.

36. Long, M.A., Katlowitz, K.A., Svirsky, M.A., Clary, R.C., Byun, T.M., Majaj, N., Oya, H., Howard, M.A., 3rd, and Greenlee, J.D. (2016). Functional Segregation of Cortical Regions Underlying Speech Timing and Articulation. Neuron 89, 1187–1193. 10.1016/j.neuron.2016.01.032.

37. Castellucci, G.A., Kovach, C.K., Howard, M.A., 3rd, Greenlee, J.D.W., and Long, M.A. (2022). A speech planning network for interactive language use. Nature 602, 117–122. 10.1038/s41586-021-04270-z.

38. Wang, R., Chen, X., Khalilian-Gourtani, A., Yu, L., Dugan, P., Friedman, D., Doyle, W., Devinsky, O., Wang, Y., and Flinker, A. (2023). Distributed feedforward and feedback cortical processing supports human speech production. Proc Natl Acad Sci U S A 120, e2300255120. 10.1073/pnas.2300255120.

39. Hage, S.R., and Nieder, A. (2016). Dual Neural Network Model for the Evolution of Speech and Language. Trends Neurosci 39, 813–829. 10.1016/j.tins.2016.10.006.

40. Hage, S.R. (2018). Dual neural network model of speech and language evolution: new insights on flexibility of vocal production systems and involvement of frontal cortex. Current Opinion in Behavioral Sciences 21, 80–87.

41. Nieder, A., and Mooney, R. (2020). The neurobiology of innate, volitional and learned vocalizations in mammals and birds. Philos Trans R Soc Lond B Biol Sci 375, 20190054. 10.1098/rstb.2019.0054.

42. Miller, C.T., Thomas, A.W., Nummela, S.U., and de la Mothe, L.A. (2015). Responses of primate frontal cortex neurons during natural vocal communication. J Neurophysiol 114, 1158–1171. 10.1152/jn.01003.2014.

43. Roy, S., Zhao, L., and Wang, X. (2016). Distinct Neural Activities in Premotor Cortex during Natural Vocal Behaviors in a New World Primate, the Common Marmoset (Callithrix jacchus). J Neurosci 36, 12168–12179. 10.1523/JNEUROSCI.1646-16.2016.

44. Zhao, L., and Wang, X. (2023). Frontal cortex activity during the production of diverse social communication calls in marmoset monkeys. Nat Commun 14, 6634. 10.1038/s41467-023-42052-5.

45. Nelson, A., Schneider, D.M., Takatoh, J., Sakurai, K., Wang, F., and Mooney, R. (2013). A circuit for motor cortical modulation of auditory cortical activity. J Neurosci 33, 14342–14353. 10.1523/JNEUROSCI.2275-13.2013.

46. Schneider, D.M., Nelson, A., and Mooney, R. (2014). A synaptic and circuit basis for corollary discharge in the auditory cortex. Nature 513, 189–194. 10.1038/nature13724.

47. Schneider, D.M., Sundararajan, J., and Mooney, R. (2018). A cortical filter that learns to suppress the acoustic consequences of movement. Nature 561, 391–395. 10.1038/s41586-018-0520-5.

48. Tsunada, J., and Eliades, S.J. (2020). Dissociation of Unit Activity and Gamma Oscillations during Vocalization in Primate Auditory Cortex. J Neurosci 40, 4158–4171. 10.1523/JNEUROSCI.2749-19.2020.

49. Granger, C.W.J. (1969). Investigating Causal Relations by Econometric Models and Cross-spectral Methods. Econometrica 37, 424–438.

50. Jacob, S.N., and Nieder, A. (2014). Complementary roles for primate frontal and parietal cortex in guarding working memory from distractor stimuli. Neuron 83, 226–237. 10.1016/j.neuron.2014.05.009.

51. Jovanovic, V., Fishbein, A.R., de la Mothe, L., Lee, K.F., and Miller, C.T. (2022). Behavioral context affects social signal representations within single primate prefrontal cortex neurons. Neuron 110, 1318–1326 e1314. 10.1016/j.neuron.2022.01.020.

52. Petrides, M., Cadoret, G., and Mackey, S. (2005). Orofacial somatomotor responses in the macaque monkey homologue of Broca’s area. Nature 435, 1235–1238. 10.1038/nature03628.

53. Petrides, M., and Pandya, D.N. (2009). Distinct parietal and temporal pathways to the homologues of Broca’s area in the monkey. PLoS Biol 7, e1000170. 10.1371/journal.pbio.1000170.

54. Kaas, J.H., and Hackett, T.A. (1998). Subdivisions of auditory cortex and levels of processing in primates. Audiol Neurootol 3, 73–85.

55. Hackett, T.A., Stepniewska, I., and Kaas, J.H. (1999). Prefrontal connections of the parabelt auditory cortex in macaque monkeys. Brain Res 817, 45–58.

56. Romanski, L.M., Bates, J.F., and Goldman-Rakic, P.S. (1999). Auditory belt and parabelt projections to the prefrontal cortex in the rhesus monkey. J Comp Neurol 403, 141–157. 10.1002/(SICI)1096-9861(19990111)403:2<141::AID-CNE1>3.0.CO;2-V.

57. de la Mothe, L.A., Blumell, S., Kajikawa, Y., and Hackett, T.A. (2006). Cortical connections of the auditory cortex in marmoset monkeys: core and medial belt regions. J Comp Neurol 496, 27–71. 10.1002/cne.20923.

58. de la Mothe, L.A., Blumell, S., Kajikawa, Y., and Hackett, T.A. (2012). Cortical connections of auditory cortex in marmoset monkeys: lateral belt and parabelt regions. Anat Rec (Hoboken) 295, 800–821. 10.1002/ar.22451.

59. Watakabe, A., Skibbe, H., Nakae, K., Abe, H., Ichinohe, N., Rachmadi, M.F., Wang, J., Takaji, M., Mizukami, H., Woodward, A., et al. (2023). Local and long-distance organization of prefrontal cortex circuits in the marmoset brain. Neuron 111, 2258–2273 e2210. 10.1016/j.neuron.2023.04.028.

60. Buzsaki, G., and Draguhn, A. (2004). Neuronal oscillations in cortical networks. Science 304, 1926–1929. 10.1126/science.1099745.

61. Buzsaki, G. (2006). Rhythms of the Brain (Oxford University Press).

62. Ford, J.M., Mathalon, D.H., Whitfield, S., Faustman, W.O., and Roth, W.T. (2002). Reduced communication between frontal and temporal lobes during talking in schizophrenia. Biol Psychiatry 51, 485–492. 10.1016/s0006-3223(01)01335-x.

63. Flinker, A., and Knight, R.T. (2016). A Cool Approach to Probing Speech Cortex. Neuron 89, 1123–1125. 10.1016/j.neuron.2016.02.039.

64. Garcia-Rosales, F., Lopez-Jury, L., Gonzalez-Palomares, E., Wetekam, J., Cabral-Calderin, Y., Kiai, A., Kossl, M., and Hechavarria, J.C. (2022). Echolocation-related reversal of information flow in a cortical vocalization network. Nat Commun 13, 3642. 10.1038/s41467-022-31230-6.

65. Hage, S.R., and Nieder, A. (2013). Single neurons in monkey prefrontal cortex encode volitional initiation of vocalizations. Nat Commun 4. 10.1038/Ncomms3409.

66. Cerkevich, C.M., Rathelot, J.A., and Strick, P.L. (2022). Cortical basis for skilled vocalization. Proc Natl Acad Sci U S A 119, e2122345119. 10.1073/pnas.2122345119.

67. Jurgens, U. (2009). The neural control of vocalization in mammals: a review. J Voice 23, 1–10. 10.1016/j.jvoice.2007.07.005.

68. Cope, T.E., Sohoglu, E., Sedley, W., Patterson, K., Jones, P.S., Wiggins, J., Dawson, C., Grube, M., Carlyon, R.P., Griffiths, T.D., et al. (2017). Evidence for causal top-down frontal contributions to predictive processes in speech perception. Nat Commun 8, 2154. 10.1038/s41467-017-01958-7.

69. Park, H., Ince, R.A., Schyns, P.G., Thut, G., and Gross, J. (2015). Frontal top-down signals increase coupling of auditory low-frequency oscillations to continuous speech in human listeners. Curr Biol 25, 1649–1653. 10.1016/j.cub.2015.04.049.

70. Park, H., Thut, G., and Gross, J. (2018). Predictive entrainment of natural speech through two fronto-motor top-down channels. Lang Cogn Neurosci 35, 739–751. 10.1080/23273798.2018.1506589.

71. Risueno-Segovia, C., and Hage, S.R. (2020). Theta Synchronization of Phonatory and Articulatory Systems in Marmoset Monkey Vocal Production. Curr Biol 30, 4276–4283 e4273. 10.1016/j.cub.2020.08.019.

72. Wilson, S.M., Saygin, A.P., Sereno, M.I., and Iacoboni, M. (2004). Listening to speech activates motor areas involved in speech production. Nat Neurosci 7, 701–702. 10.1038/nn1263.

73. Pickering, M.J., and Garrod, S. (2007). Do people use language production to make predictions during comprehension? Trends Cogn Sci 11, 105–110. 10.1016/j.tics.2006.12.002.

74. Cheung, C., Hamiton, L.S., Johnson, K., and Chang, E.F. (2016). The auditory representation of speech sounds in human motor cortex. Elife 5. 10.7554/eLife.12577.

75. Pomberger, T., Risueno-Segovia, C., Loschner, J., and Hage, S.R. (2018). Precise Motor Control Enables Rapid Flexibility in Vocal Behavior of Marmoset Monkeys. Curr Biol 28, 788-+. 10.1016/j.cub.2018.01.070.

76. Nummela, S.U., Jovanovic, V., de la Mothe, L., and Miller, C.T. (2017). Social Context-Dependent Activity in Marmoset Frontal Cortex Populations during Natural Conversations. J Neurosci 37, 7036–7047. 10.1523/JNEUROSCI.0702-17.2017.

77. Eliades, S.J., and Wang, X. (2008). Chronic multi-electrode neural recording in free-roaming monkeys. J Neurosci Methods 172, 201–214. 10.1016/j.jneumeth.2008.04.029.

78. Agamaite, J.A., Chang, C.J., Osmanski, M.S., and Wang, X.Q. (2015). A quantitative acoustic analysis of the vocal repertoire of the common marmoset (Callithrix jacchus). J Acoust Soc Am 138, 2906–2928. 10.1121/1.4934268.

79. Brovelli, A., Ding, M., Ledberg, A., Chen, Y., Nakamura, R., and Bressler, S.L. (2004). Beta oscillations in a large-scale sensorimotor cortical network: directional influences revealed by Granger causality. Proc Natl Acad Sci U S A 101, 9849–9854.

80. Jacob, S.N., Hahnke, D., and Nieder, A. (2018). Structuring of Abstract Working Memory Content by Fronto-parietal Synchrony in Primate Cortex. Neuron 99, 588–597 e585. 10.1016/j.neuron.2018.07.025.

81. Ninomiya, T., Noritake, A., Kobayashi, K., and Isoda, M. (2020). A causal role for frontal cortico-cortical coordination in social action monitoring. Nat Commun 11, 5233. 10.1038/s41467-020-19026-y.

82. Noritake, A., Ninomiya, T., Kobayashi, K., and Isoda, M. (2023). Chemogenetic dissection of a prefrontal-hypothalamic circuit for socially subjective reward valuation in macaques. Nat Commun 14, 4372. 10.1038/s41467-023-40143-x.

83. Epple, G. (1968). Comparative Studies on Vocalization in Marmoset Monkeys (Hapalidae). Folia Primatol 8, 1-&. Doi 10.1159/000155129.

84. Bezerra, B.M., and Souto, A. (2008). Structure and usage of the vocal repertoire of Callithrix jacchus. Int J Primatol 29, 671–701. 10.1007/s10764-008-9250-0.

85. Oostenveld, R., Fries, P., Maris, E., and Schoffelen, J.M. (2011). FieldTrip: Open source software for advanced analysis of MEG, EEG, and invasive electrophysiological data. Comput Intell Neurosci 2011, 156869. 10.1155/2011/156869.

86. Pesaran, B., Vinck, M., Einevoll, G.T., Sirota, A., Fries, P., Siegel, M., Truccolo, W., Schroeder, C.E., and Srinivasan, R. (2018). Investigating large-scale brain dynamics using field potential recordings: analysis and interpretation. Nat Neurosci 21, 903–919. 10.1038/s41593-018-0171-8.

